# Exploring large protein sequence space through homology- and representation-based hierarchical clustering

**DOI:** 10.1101/2024.11.13.623527

**Authors:** JZ Chen, B Gall, N Tokuriki, CJ Jackson

## Abstract

Exploration of protein sequence space can offer insight into protein sequence-function relationships, benefitting both basic science and industrial applications. The use of sequence similarity networks (SSNs) is a standard method for exploring large sequence datasets, but is currently limited when scaling to very large datasets and when viewing more than one level (hierarchy) of homology. Here, we present a sequence analysis pipeline with a number of innovations that address some limitations of traditional SSNs. First, we develop a hierarchical visualization approach that captures the full range of homologies across protein superfamilies. Second, we leverage representations embedded by protein language models as an alternative homology metric to the basic local alignment search tool (BLAST), showing that they produce comparable results when identifying isofunctional protein families. Finally, we demonstrate that unbiased representative sampling of sequences from genetic neighborhoods can be achieved through the use of hidden Markov models (HMMs) or vector representations. The utility of these methods is exemplified by updating the sequence-function analysis of the FMN/F_420_-binding split barrel superfamily and improving phylogenetic analyses. We provide our sequence exploration pipeline as publicly available code (ProteinClusterTools) and show it to be scalable to large datasets (∼300k sequences) using desktop computers.

## Intro

Knowledge of protein sequence-function relationships is important in biology and protein engineering as it allows for a better understanding of the factors that underlie protein functions, and to predict or design biological functions based on the amino acid sequence. A rich source of sequence-function information lies in protein sequence space — the collection of all known protein sequences and their functions. From a theoretical perspective, protein sequence space can be viewed at different levels of organization that captures evolutionary history and function (**Fig. 1a**) ^1–6^. At the broadest level, highly divergent proteins form evolutionarily distinct ‘superfamilies’, often featuring a conserved fold, and are host to diverse functions across different host organisms. Superfamilies can then be divided into various ‘families’ (isofunctional homologs), often distinguished by their sequence homology. In practice, gleaning insight into sequence space requires researchers to empirically determine the functional boundaries of different protein families by dividing the sequence space into closely and distantly related sequence “neighborhoods”.

**Figure 1.**
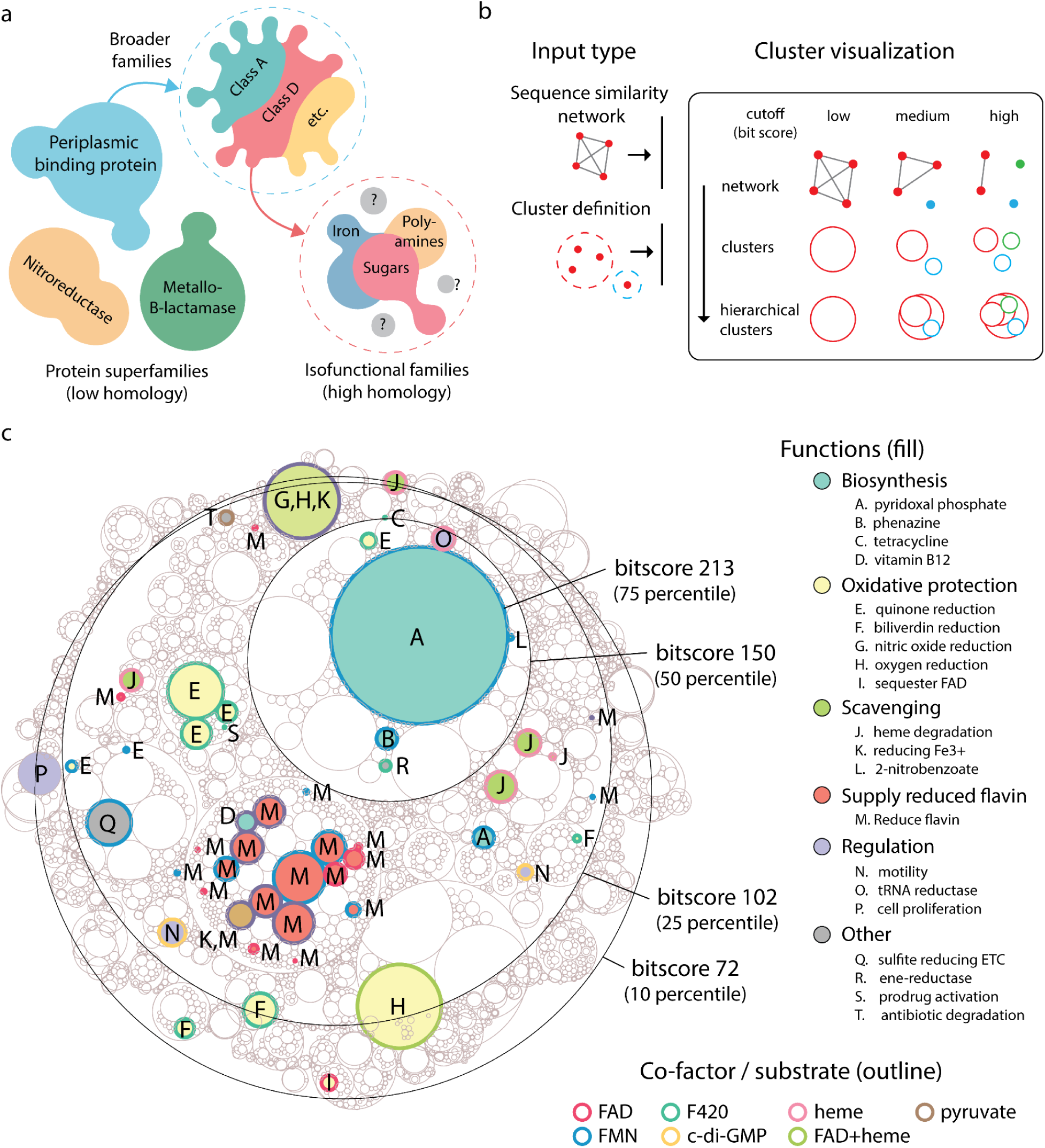
Protein sequence space organization and workflow for exploration. (**a**) Schematic overview of protein sequence space at different levels of homology. (**b**) Schematic detailing the visualization logic of a hierarchical cluster plot, with entry points for different types of sequence neighborhood data. (**c**) SSN of the FMN-binding split barrel superfamily visualized as a hierarchical cluster plot. Functions and co-factors are annotated at the highest bitscore level (213). Each cut-off is highlighted by an example neighborhood at that cut-off. Neighborhoods at bitscore 72 that did not have any annotated functions were hidden.

Thanks to advances in deep sequencing and genomics, our view of sequence space is ever expanding with the growth in genomic and metagenomic data. However, the number of sequences with empirically determined functions (*i.e.*, that have been characterized) within this sequence space remains very small. At the time of writing, there are over 245 million protein sequences in the UniProtKB database, yet only ∼0.2% (∼0.57 million) of those sequences are matched to curated SwissProt functional annotations. By identifying sequence neighborhoods, the sparse collection of proteins with known functions can be used to infer functions of closely related sequences in the same neighborhood and expand our knowledge of functional sequence space ^6–9^. At the same time, neighborhoods with no associated functions will stand out as gaps in our knowledge, representing potentially exciting opportunities for exploring new functions ^6,10,11^.

The sequence similarity network (SSN) is the most common approach for exploring neighborhoods in sequence space ^12–14^. Sequences are represented as nodes on a graph and are connected by edges to other nodes with which they share similarity above a given cut-off ^13^. By applying layout algorithms, sequence neighborhoods emerge through the distinct clustering of closely related sequences. Identifying the ideal similarity cut-off requires manual optimization and can reveal neighborhoods of sequences that define functional families^12^; i.e., the cut-off should be sufficiently high to exclude different functions yet not too strict such that only extremely similar sequences remain connected. Despite their utility, SSNs currently suffer limits in scalability as large networks (e.g., >3M edges) are hard to visualize, often requiring reducing the dataset and discarding connections. Furthermore, SSNs can only visualize a single cut-off at a time which limits the ability to accommodate different isofunctional families (which often group at different cut-offs) in the same overview of the superfamily. A recent structure-based sequence space exploration approach (ProteinCartography) offers an alternative to traditional SSNs, using dimensionality reduction (t-SNE and UMAP) to visualize the protein space and removing the need for edges; though the Leiden clustering used to define sequence groupings only provides a single level of separation (i.e., a fixed single cut-off) ^15^.

In this work, we build on the traditional pipeline of protein sequence space exploration to improve and complement the existing tools and demonstrate the utility of these new approaches on the FMN/F_420_-binding split barrel superfamily (∼300k sequences)^16^. We introduce a ‘hierarchical cluster plot’ that provides succinct, interactive overviews of sequence neighborhoods across different levels of homology. This new visualization method also addresses the lack of scalability in conventional SSN visualization, bypassing the depiction of edges. We test alternatives to the traditional homology based neighborhoods in SSNs by using sequence vector representations from protein language models (pLM)^17–19^ for sequence space exploration, and find them to perform comparably to the homology based method in neighborhood assignment, with the advantage of comparing full sequences without alignments. We then explore methods of selecting the most representative sequences from sequence neighborhoods to allow for systematic and unbiased analysis of sequence space. Finally, we demonstrate the utility in using sequence space information to construct a phylogenetic tree of the FMN/F_420_-binding split barrel superfamily, showing improved sequence coverage and replicability of tree topology, compared to the conventional approach (e.g., BLAST of a few sequences selected on the basis of available structures). We combine these innovations into an analysis pipeline (ProteinClusterTools) and have made the code available to facilitate ease-of-use in analyzing large protein superfamilies.

## Results

### Exploring the FMN-binding split barrel superfamily with hierarchical information

Sequence space can be thought of as a hierarchical organization that can be resolved at different levels of homology, from evolutionarily distinct superfamilies to more closely related isofunctional families (**Fig. 1a**). To account for this hierarchical nature and improve the efficiency for analyzing large datasets, we developed the “hierarchical cluster plot”, a cluster visualization method that captures this organization to aid in visualization of sequence neighborhoods. First, isolated clusters (no connecting edges to another cluster) are determined from an underlying SSN at each cut-off (bitscore) in the sequence network. Then, clusters of lower cut-offs (broader homology) act as parent “neighborhoods” for clusters that further separate at higher cut-offs (higher homology) (**Fig. 1b**).

To demonstrate the utility of this method, we created an SSN of the FMN/F_420_-binding split barrel superfamily (also known as flavin/deazaflavin oxidoreductases, or FDORs) which was previously analyzed using ∼2k sequences ^16^. Here, we extend the SSN to the entire FMN-binding split barrel superfamily from UniProt (IPR012349 and IPR037119), expanding the dataset by ∼150x (∼300K non-redundant sequences, ∼280M edges). Homology is measured by the bitscore (BLOSUM62 matrix), calculated from pairwise local alignments using MMSeqs. To evenly select cut-offs for visualization, we sample the distribution of all bitscores and identify the bitscores corresponding to the 10, 25, 50, and 75 percentiles of the distribution (bitscores of 72, 102, 150, 213) and overlay each cut-off in a hierarchical cluster plot (**Fig. 1c**). In contrast to visualizing SSNs directly as networks, the hierarchical cluster plot approach avoids the rendering of individual nodes and edges, making visualizations considerably more efficient. When visualizing the SSN directly as a network in Cytoscape, we find a linear scaling of memory usage and an exponential increase in time taken to produce the force directed layout (required to see clusters visually) (**Supp. Fig. 1**, **Supp. Table 1**). Notably, the layout time presents a major time bottleneck for networks with >3M edges, and the cut-offs analyzed in the hierarchical cluster plot would be impossible using conventional networks (**Supp. Table 2**).

The superfamily was annotated for function and co-factor binding using available data from UniProt and the PDB (**Supplementary Data 1**). We define isofunctional families (“families” from here onwards) as clusters with only one annotated function at the highest cut-off, while a neighborhood is the common parent cluster to a group of child clusters (*i.e.*, the clusters are closer to each other than to other neighborhoods). Most functions in the superfamily fall within five major functional groups: biosynthesis, oxidative protection, nutrient scavenging, supplying reduced flavins and regulation. At the 75 percentile cut-off (bitscore 213), most clusters are isofunctional. The most commonly annotated function is for proteins that supply reduced flavin to a downstream enzyme, usually as two-component systems^20,21^. Seventeen different families (2-2025 sequences) occupy the same neighborhood, possibly forming a single larger family. Five other small families (>=24 sequences) emerge separately throughout the superfamily. All proteins that supply flavins bind FAD or FMN. The largest biosynthetic family is for pyridoxal phosphate, formed by PNP oxidases (PNPOx)^16,22,23^ with ∼24k sequences, with a single, separate family in a separate neighborhood (330 sequences). The family for phenazine synthesis ^24^ (327 sequences) shares a neighborhood with the large PNPOx family, while tetracycline^25^ and vitamin B12 ^26^ biosynthesis are in separate neighborhoods. All biosynthetic families use either FAD or FMN, save for tetracycline, which uses the deazaflavin cofactor, F_420_. There are a number of families involved in oxidative protection. Quinone reductases^16,27,28^ are the most common, with three families (∼300-2400 sequences each) utilizing F_420_ from the same neighborhood, suggesting that they belong to the same larger family. Another F_420_ utilizing quinone reductase family is in the large PNPOx neighborhood, while 2 smaller families(<=79 sequences) of FMN utilizing quinone reductases arise in a separate neighborhood. Biliverdin reduction ^16,29^ is reported in three separate families each in its own neighborhood (32-863 sequences), all of which utilize F_420_ cofactors. There are two clusters with enzymes involved in removing O_2_ or NO gases ^30–32^, with one mixed cluster including both functions, and a separate family that removes only O_2_. Finally, one family has been reported to sequester FAD to prevent oxidation and ROS generation^33^. Heme degradation ^16,34,35^ is the most common function in proteins associated with nutrient scavenging, with three families in the same neighborhood (22-727 sequences). Two other heme degradation families emerged separately. There are two families that reduce Fe^3+^ to release iron from siderophores ^36,37^, and one small family catabolizing 2-nitrobenzoate^38^ in the same neighborhood as the large PNPOx family. A number of families perform non-enzymatic regulatory functions: two families bind cyclic di-GMP to regulate motility ^39,40^, while one family regulates glutamyl-tRNA reductase^41^. Another family has been reported to be involved in regulating cell proliferation^42^. The final observation that can be drawn from this network is that a vast amount of the known sequence space in this family remains uncharacterized, including several very large clusters.

The use of the hierarchical cluster plot in this analysis demonstrates a number of benefits. Different parts of sequence space can have different cut-offs for separating into isofunctional families, where some families can be isolated at lower homologies while others only separate at higher homology levels. A hierarchical view can accommodate cut-offs for all families not possible with just one fixed cut-off. The hierarchical structure also provides information across cut-offs on the diverging relationship of families from distant relatives to close homologs, and may help highlight whether families may have arisen divergently (emerging from the same neighborhood), or convergently (different neighborhoods across the superfamily). Clusters of interest can also be selected for higher resolution separation at higher cut-offs, or to extract a more manageable subset of the SSN for direct viewing. In addition, the cluster based approach is compatible with any set of cluster definitions, regardless of how the clusters were generated. Thus, the hierarchical cluster plot is a scalable and flexible visualization method that succinctly summarizes broad organization in sequence space.

### Use of vector representations in sequence neighborhood detection

With the advent of new pLMs in the study and engineering of proteins, we explore the potential of pLM embeddings as a new homology metric for exploring sequence neighborhoods. In conventional SSNs, pairs of sequences are compared using alignments and their homology is measured by scoring using a substitution matrix^13^. The most commonly used substitution matrix, BLOSUM62, captures substitution behaviors in conserved blocks of distantly related proteins^43^. However, the gaps in the alignments use heuristic scores for insertions/deletions set by the user, without exact knowledge of insertion/deletion likelihoods. Furthermore, SSNs using local alignments will only evaluate homology based on the aligned region, and exclude other non-aligned regions such as extra domains. More recently, advancement in machine learning has led to the creation of protein language models (pLMs), which are trained on all known proteins in order to recognize their sequence patterns^17–19^. The patterns learned by pLMs can be extracted, by using the pLM to encode any given protein sequence as a numerical vector. The exact vector has no meaning, but differences between vectors correlate with differences in sequence identity (**Supp. Fig. 2**) and can be used to recognize sequence-function relationships^17^. Hence, we can also utilize pLM sequence representations (from the pLM ESM1b in our case) to analyze protein sequence space, which can be applied to clustering algorithms in an alignment free way and while comparing the full sequences.

We apply bisecting K-means (**Fig. 2a**) or hierarchical clustering (**Fig. 2b**) on the vector representations (methods). Bisecting K-means is a divisive clustering algorithm that starts with one cluster with all sequences, then performs divisions until the desired number of clusters (K parameter) is reached; at each step the cluster with the largest within cluster difference is chosen and split into 2 smaller clusters. In contrast, hierarchical clustering is an agglomerative clustering algorithm that starts with all sequences as separate clusters, then joins the closest clusters in pairs until all are in the same cluster. Hierarchical clustering can also produce a tree structure that tracks the stepwise separation of all sequences and clusters, allowing the researcher to decide on the best neighborhood definitions (clades in tree) at any depth within the tree structure (**Fig. 2c**). By clustering the sequence space of the FMN-binding split barrel binding superfamily as vector representations into 5, 10, 100 and 1000 clusters, we can produce a hierarchical plot analogous to the one yielded by the conventional homology method in **Fig. 1c**. Qualitatively, the vector representations yield similarities to the homology based neighborhoods. Enzymes supplying reduced flavins to other enzymes (M) form large neighborhoods together and share neighborhoods with vitamin B12 synthesis (D) and Fe^3+^ reduction (K). The PNPOx enzymes (A) also group into one large neighborhood, with a smaller family that groups separately. The F_420_ utilizing quinone reductases (E, green outline) form a separate neighborhood from the FMN utilizing family (E, blue or purple outline). One difference is that the vector representation shows distinct separation of cofactors, with most F420 enzymes consistently grouped into the same parent neighborhoods (K or clusters=5), while no distinct grouping of F420 appears in the homology based methods.

**Figure 2.**
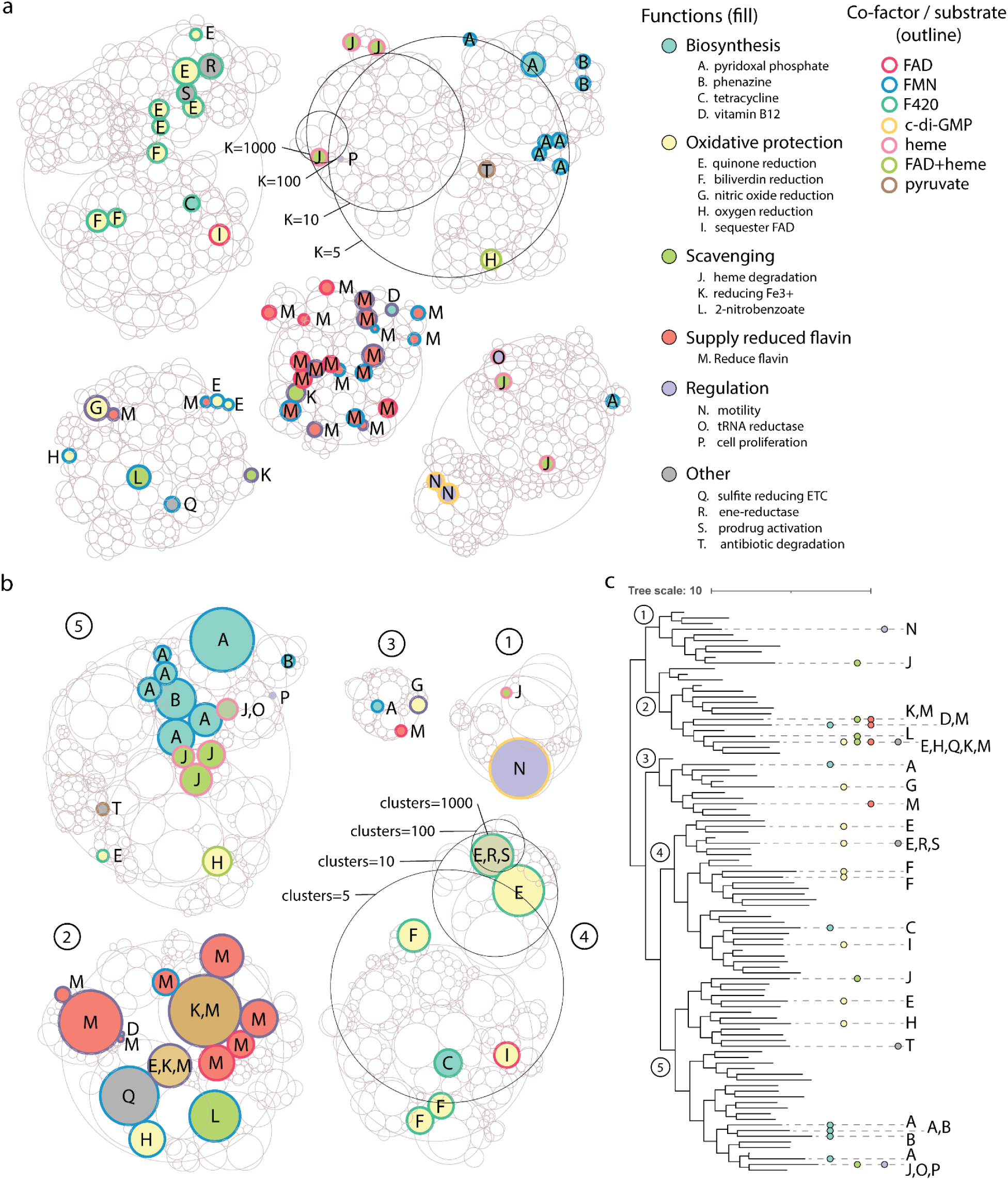
Clustering patterns of vector representation based clusterings. (**a**) Hierarchical cluster plot of bisecting K-means clustering with euclidean distance at K=5,10,100 and 1000. (**b**) Hierarchical cluster plot of hierarchical clustering with weighted linkage function and cosine similarity as distance, extracted at clusters=5,10,100 and 1000. Clusters are numbered at clusters=5. (**c**) Tree structure of hierarchical clustering shown in **b** (matching numbered clades), with branches collapsed to the 100 clusters cut-off. Branch lengths represent the longest distance to any single sequence in the collapsed clades. Functions for each collapsed cluster are highlighted to the right. Tree visualized with interactive tree of life (ITOL).

To have a quantitative understanding of clustering similarities, we calculate clustering agreement using the adjusted rand-index (ARI) and adjusted mutual information (AMI). For both metrics, 0 means the cluster assignments are random or independent, while 1 means perfect agreement. We compare cluster definitions generated at a range of cut-off for each pair of methods (homology vs K-means, homology vs hierarchical, K-means vs hierarchical) and find cluster assignments to be generally similar between methods (**Supp. Fig. 3**). Between homology and vector methods ARI can range up to 0.27 (vs K-means) to 0.32 (vs hierarchical), while AMI can range up to 0.63 (vs K-means) to 0.66 (vs hierarchical). Within the vector methods the agreement is higher, with maximum ARI of 0.53 and maximum AMI of 0.73 between K-means and hierarchical clustering. The exact agreement is cut-off dependent, likely due to the effect of differing cluster count and cluster sizes on the agreement metrics (*e.g.*, K-means and hierarchical agree better at more similar cluster counts). Hence, we have shown that vector representations can also be used to identify sequence neighborhoods in a superfamily, with comparable broad level trends and similarities in exact clustering to those produced by homology.

### Properties of different clustering methods

The key to extracting information from protein superfamilies is to collect groups of related sequences rather than treat all data as individual sequences. To better understand the utility of our clustering, we examine the properties of the neighborhoods produced by each of the clustering methods on the FMN-binding split barrel binding superfamily (for clusters with N>=5). We analyze the cluster definitions at the previously presented cut-offs, where lower cut-offs were able to show broad trends (grouping of similar functions) and the highest cut-offs were able to reveal similar isofunctional neighborhoods (**Fig. 1b** and **Fig. 2**).

We examine the size distribution of clusters produced at each cut-off (**Fig. 3a**). We find that the homology based approach produces small clusters across all cut-offs, while the vector approaches produce a range of cluster sizes depending on the cut-off. Within vector approaches, the K-means approach has narrower distributions of cluster sizes compared to hierarchical clustering; the evenly sized K-means clusters are apparent in the hierarchical cluster plots (**Fig. 2**). To consider the consistency in sequence membership, we explore the distribution of pairwise identities within each cluster of the different methods (**Fig. 3b, Methods**). For all methods, the baseline sequence identity is ∼30-40% when comparing random pairs from different clusters at the highest degree of separation. Within methods, the sequence identity within neighborhoods gradually increases as the neighborhoods become more numerous (and hence smaller). The homology based approach converges to more similar sequences within clusters compared to the vector methods, despite both arriving at similar final clusterings. Finally, we examine the taxonomic diversity of each neighborhood and find a trend similar to the progression of neighborhood sizes (**Fig. 3c**). As expected, taxonomic diversity is highest at the lowest neighborhood count (largest clusters), and gradually decreases. Again, we see the homology approach produces clusters with lower taxonomic diversity through all cut-offs. Meanwhile, the K-means and hierarchical clustering result in having a higher starting diversity at most lower K cut-offs, but eventually converging to a smaller number of taxa at higher cut-offs.

**Figure 3.**
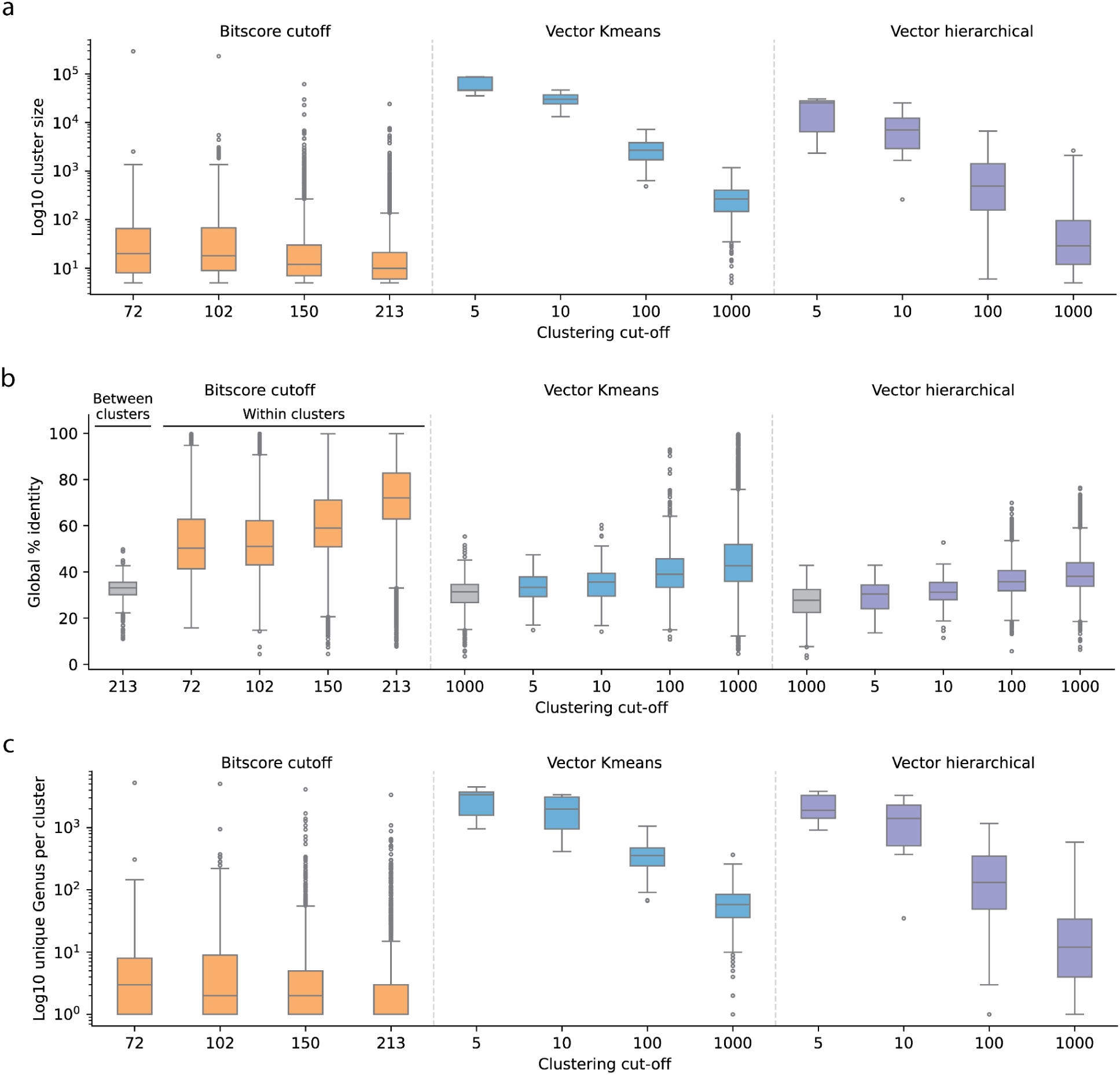
Comparison of behaviors across different clustering methods. (**a**) The distribution of cluster sizes when the sequence space is split into clusters at a given cut-off. (**b**) The distribution of global sequence identity between sequences of separate clusters (gray) and between sequences within the same cluster (colored). (**c**) The distribution of taxonomic diversity (number of unique genuses) within clusters at each clustering cut-off. For **a-c**, all clusters analyzed have a minimum of 5 sequences.

These observations are consistent with the expected tendencies of each clustering method; specifically, that the homology approach tends to produce smaller clusters with more similar sequences and lower taxonomic diversity than the K-means or hierarchical clustering. For instance, in the clustering approach of identifying isolated clusters above a homology cut-off, even a single connection between two otherwise separated clusters will convert it into a large cluster, and hence clusters that do separate will likely be smaller and more self-similar. In the case of K-means, the evenness in the progression and distribution spread of cluster size and taxonomic range is consistent with the tendency of bisecting K-means, which prioritizes dividing clusters that have the most unevenness within the cluster. Visually, the hierarchical clustering appears to be an intermediate between the homology and K-means approach, with more separation than the homology approach, but less clusters than K-means (**Fig. 1b** vs **Fig. 2**). It is interesting that despite overall similar clustering outcomes, the methods produce neighborhoods that are quite different in terms of size and sequence identity. We infer this to mean that detection of isofunctional families, where the data labels are very sparse and widely distributed, would not significantly be affected by such differences in neighborhood properties. However, if the researcher is interested in properties of whole clusters, the choice of method could be important. For example, when desiring sequences that are very similar in terms of sequence identity, one may favor the homology method, while K-means could produce the most evenly sized neighborhoods that still adhere to functional separation.

### Extracting cluster representatives

Once a sequence neighborhood has been identified, it is often necessary to reduce the data further for downstream processes, such as experimental characterization or phylogenetics analysis. We present two different methods for selecting representatives given a cluster of sequences (**Fig. 4a**). One method relies on the construction of a multiple sequence alignment of all member sequences within the cluster. The MSA is used to build a sequence profile HMM, which can be used to test all member sequences similarity to the common profile. The best representative is the member with the closest similarity to the HMM. In the second method, the vector representations of all member sequences are averaged to get the vector centroid of the cluster. Then the distance of all members to the centroid is evaluated, and the representative is the sequence closest to the centroid.

**Figure 4.**
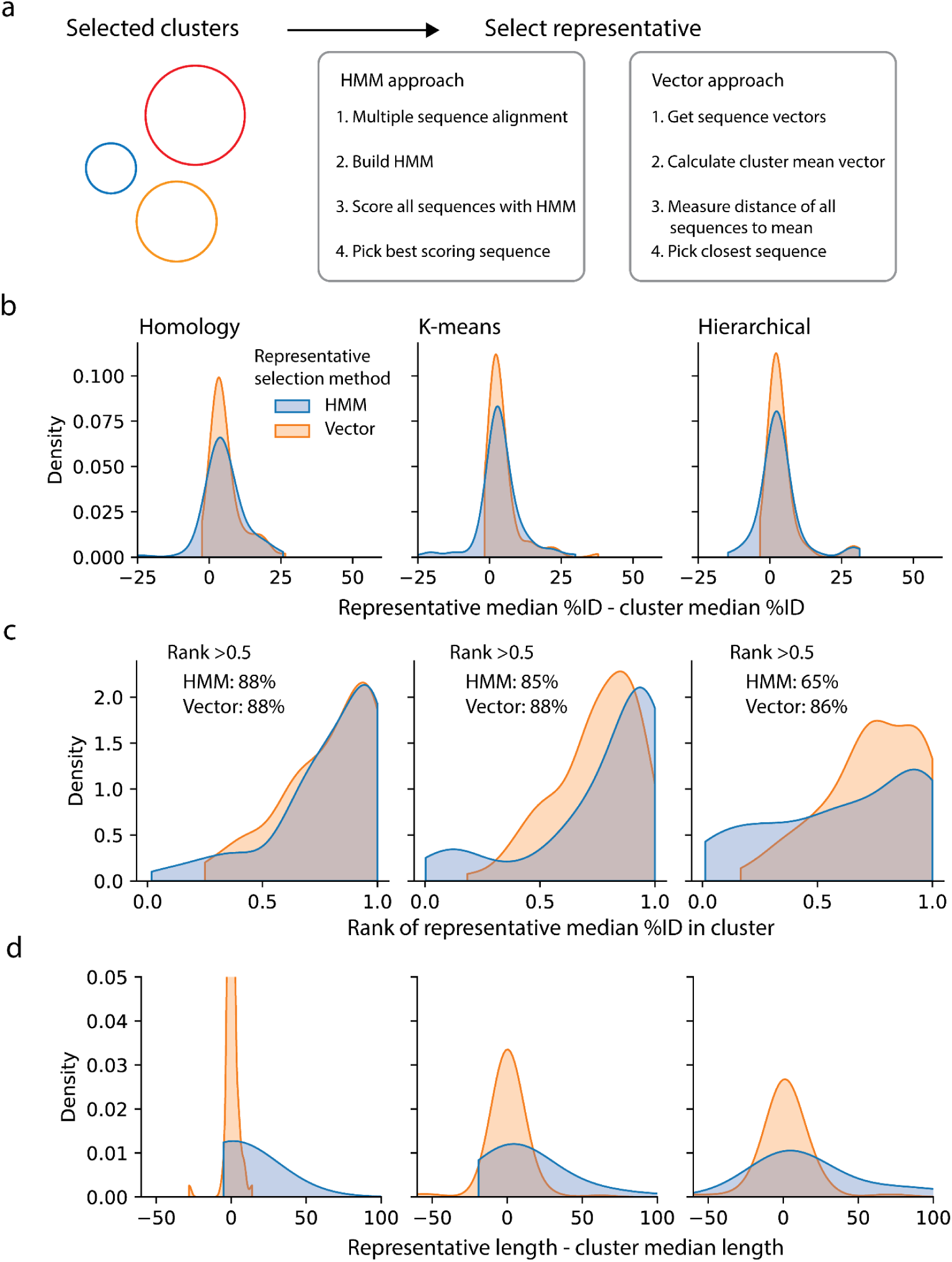
Methods for selecting representative sequences from sequence clusters. (**a**) Schematic of two approaches to selecting representatives from sequence clusters, either using HMMs or the vector representation of sequences. (**b**) The distribution of differences in sequence identity, between the %ID of the selected representative to all cluster members and the median %ID of random pairs of cluster members. (**c**) Distributions of the representatives’ percentile rank within each cluster, when comparing median %ID to the cluster calculated for all members of the cluster. (**d**) The distribution of differences in sequence length, between the selected representative and the median cluster length, plotted for each representative selection method and each clustering method.

We applied both representative selection approaches to each clustering method, then assessed how well the selected representative represents the cluster. We evaluated the highest cut-off of each clustering method (*i.e.*, where most neighborhoods are already isofunctional) and randomly selected 100 clusters with between 5-1000 sequences. To compare the representatives against random selection of sequences, we first calculated the median sequence identity of all cluster members to the representative, then compared this to the median random pairwise sequence identities within the cluster (**Fig. 4b**). We found ∼79-93% representatives are closer to all other cluster members in terms of sequence identity than random pairwise identities. To determine how well the representative represents the cluster compared to other members, we computed the median sequence %ID of each member to all other members in the cluster, then ranked all median sequence %IDs within the cluster (**Fig. 4c**). We found that both HMM and vector representatives tended to be more similar to the cluster than most other members, with between 65-88% of representatives being more similar to the cluster than half of the cluster (rank > 0.5). Finally, to examine if sequence length is properly represented, we calculated the difference in length of the chosen representative sequence to the median sequence length within the cluster (**Fig. 4d**). Again, we found the representative to be a good match to the cluster. The representative is typically within ∼50 amino acids of the cluster median length, meaning the difference in most cases is smaller than a typical protein domain^44^, even small domains like DNA binding domains. We note the HMM method exhibits a bias toward higher lengths and greater spread in length differences compared to the vector method. Overall, both methods tend to produce representatives that represent the members of the clusters well.

### Leveraging diverse sequence neighborhoods for phylogenetic trees

One application that would benefit from a complete overview of sequence space is the sampling of sequences for phylogenetic trees. Typically, phylogenies of protein families are constructed by taking representative members of the family (*e.g.*, having known function or structure) and conducting homology searches such as BLAST to retrieve sequences in the neighborhood. Such a method is simple and ensures key sequences of interest are covered, but may not fully cover the sequence space between different distinct subfamilies. A previous study constructed a phylogenetic tree of the FMN-binding split barrel superfamily (also referred to as β-roll F_420_-dependent enzymes) using such an approach ^45^. To assess the sequence space coverage, we replicated the sampling method by taking the top 500 BLAST hits of 18 structural representatives (PDB ID 2FQH was excluded as it was a different fold), and reduced the redundancy of sequences to 65% sequence identity. Although the previous study used 80% identity to reduce redundancy, we still obtained a greater number of sequence representatives for the tree after cleaning (321 now vs 105 previously). Despite the growth in sequences, we find the coverage of the BLAST method to be somewhat sparse in view of the full sequence space (**Fig. 5a**). We then devised a sampling method using our cluster tools to leverage the full sequence space using the homology based SSN. We sample one representative from each neighborhood that’s between 100-1000 sequences starting at a bitscore of 150 (only parent neighborhoods at 102 bitscore that contain any BLAST representative are considered). Neighborhoods with more than 1000 sequences are further separated into smaller neighborhoods in steps of 10 bitscore, up to 400. This resulted in a comparable number of representatives for the tree to the BLAST method (364 including the 18 structural representatives), but the coverage of the full sequence space is greatly improved (**Fig. 5a**).

**Figure 5.**
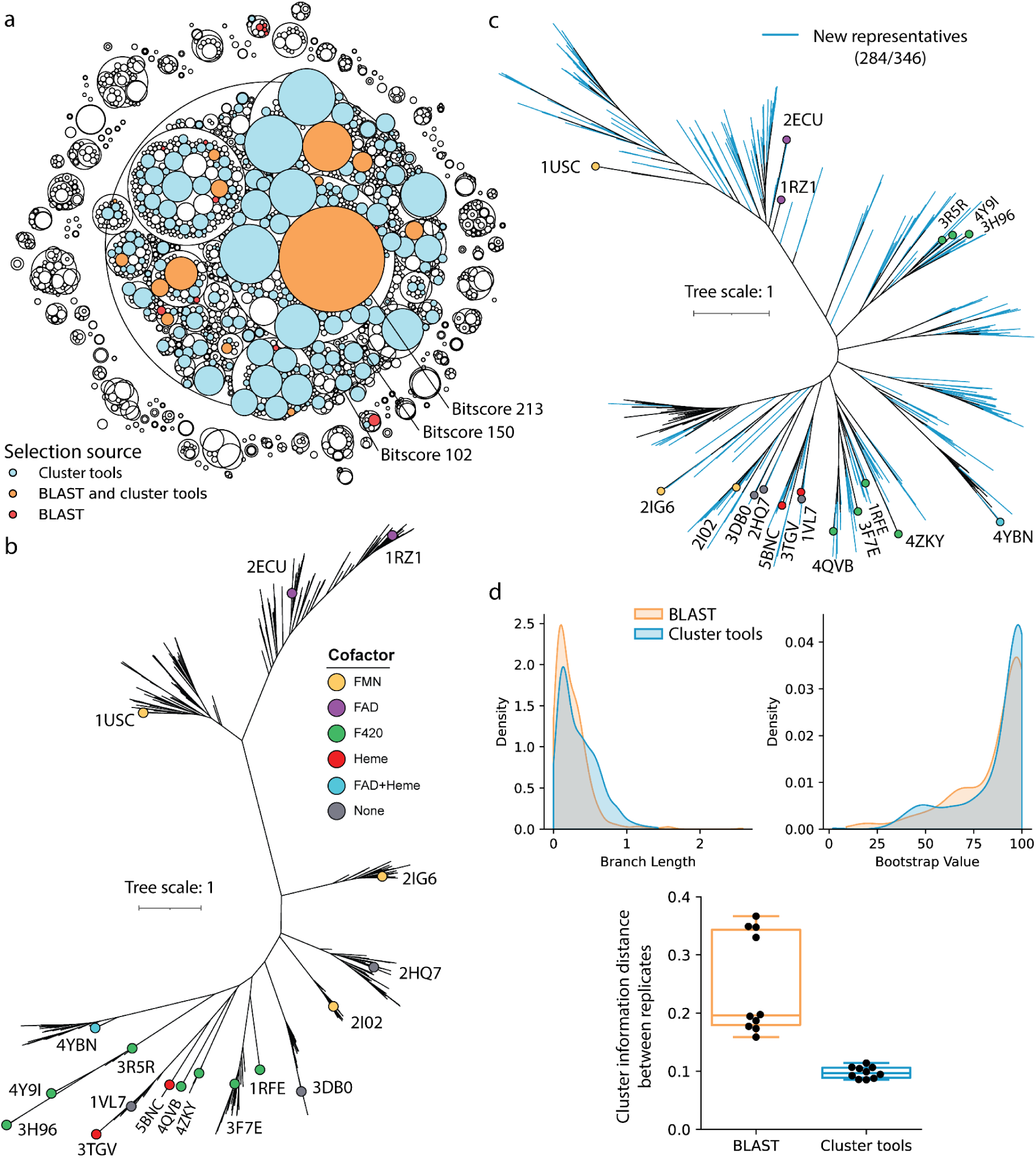
Phylogenetics of FMN-binding split barrel superfamily using diverse sequence coverage. (**a**) Hierarchical cluster plot of the FMN-binding split barrel superfamily at bitscore cut-offs of 102, 150 and 213, highlighted by the sequence coverage of different sequence sampling methods. (**b**) Phylogenetic tree constructed from BLAST sampled sequences (321 leaves). The 18 structural representatives and their cofactors are highlighted on the tree. Fully labeled tree in **Supplementary figure 4**. (**c**) Phylogenetic tree constructed from evenly sampling clusters in the SSN (364). The 18 structural representatives and cofactors are labeled. Blue branches indicate sequences from clusters that were not sampled in the BLAST method. Fully labeled tree in **Supplementary figure 5**. (**d**) Statistics for trees generated using both methods, including the distribution of all branch lengths, bootstrap values, and tree distance between different replicates (5 replicates each, N=10 pairwise distances).

To examine the effects of the different sampling methods, we construct maximum likelihood phylogenetic trees from each set of sampled sequences. In terms of topology, the BLAST based tree (**Fig. 5b**) exhibits clades of closely related sequences that are separated by very long branches. In contrast, the cluster tools based tree (**Fig. 5c**) shows a more evenly spread topology in terms of clades and connecting branch lengths. This is captured by the distribution of branch lengths in each tree (**Fig. 5d**). Both trees share general similarities, such as the grouping of F_420_ utilizing quinone reductases (PDB IDs: 3R5R, 3H96, 4Y9I, a.k.a. FDOR-A ^16^) and FAD utilizing flavin reductases (PDB IDs: 2ECU, 1RZ1). However, with the addition of more diverse sequence data using cluster tools, there is more distinct separation between the heme binding proteins (PDB IDs: 3TGV, 5BNC) and the F420 binding proteins (PDB IDs: 1RFE, 3F7E, 4QVB, 4ZKY, a.k.a FDOR-B ^16^) (**Fig. 5c**), which formed an intermixed clade in the BLAST based tree (**Fig. 5b**). In terms of reliability we find the bootstrap values to be comparable between both methods, but the use of more diverse representatives lead to more consistent tree topology between replicates as shown by the lower cluster information distance^46^ (**Fig. 5d**). In addition, the cluster tools based sampling greatly expands the sequence space covered, where 82% of sequences represent clusters not covered by the BLAST method despite both sampling a comparable number of sequences. As a result, a number of new clades can be observed that were not present in the BLAST based tree (**Fig. 5c**). Hence, we find that leveraging the full sequence space for phylogenetic tree construction can improve the coverage and replicability of tree topology, while achieving similar bootstrap values.

### A pipeline for sequence space exploration

We incorporate our methods into a pipeline with the aim of streamlining sequence space exploration. The typical workflow for sequence space exploration is shown in **Fig. 6**. The researcher first identifies the target protein family or superfamily they wish to study, and collects the protein sequences from a public database^47^. Next, an SSN is constructed using an all by all BLAST to visualize sequence neighborhoods in the form of clusters, and can be annotated with various features available to the user (most commonly, the proteins’ function). By itself, the full SSN can provide a good overview of the protein family for broad level bioinformatics. The sequence neighborhoods can be further broken down into smaller neighborhoods at cut-offs of higher homology to provide higher resolution separation between sequences. Once a desirable cut-off is chosen, the user can select neighborhoods of interest and choose representative sequences for further downstream characterization.

**Figure 6.**
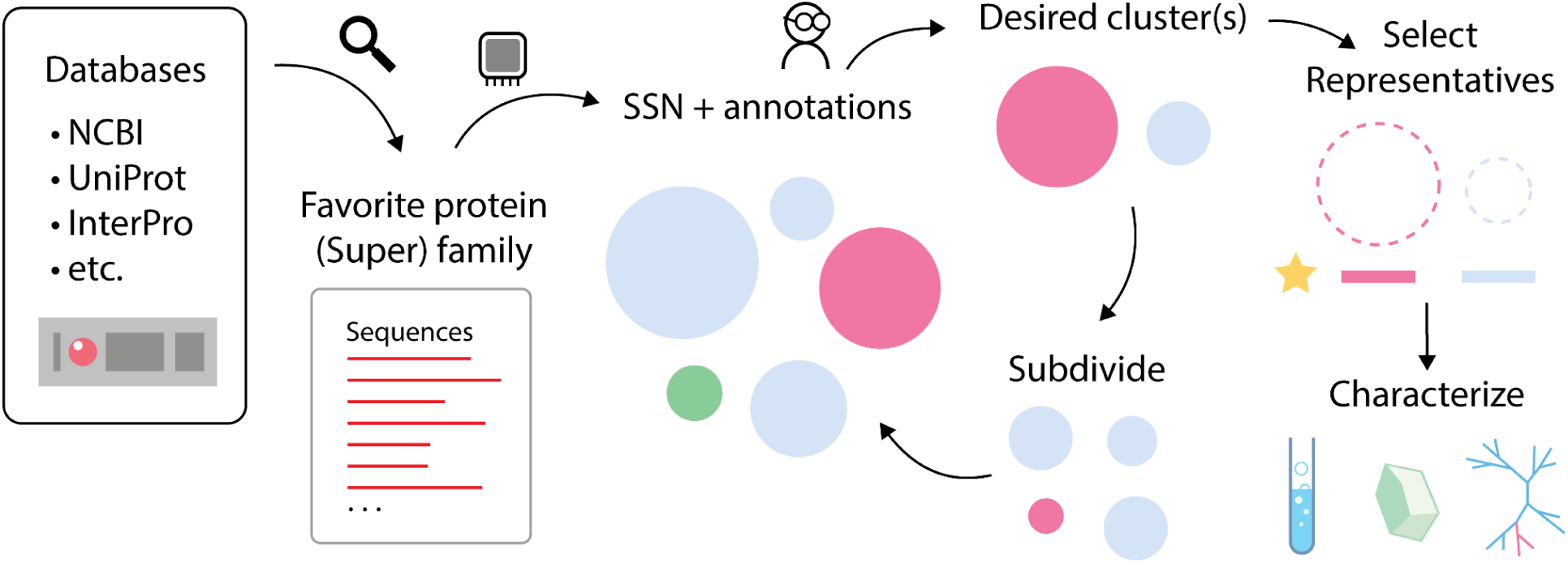
Workflow for sequence space exploration. General workflow for the pipeline aimed at examining protein sequence space, identifying sequence neighborhoods, and selecting representatives for downstream characterization.

The pipeline implements all neighborhood detection methods described here (homolog, K-means and hierarchical clustering), allowing the user to conduct all analysis starting from just a single fasta file of sequences. With regards to the homology method, we implement a new method for detecting connected components aimed at scalability for large network files that are too large to load into RAM. The neighborhood definitions can then be directly visualized as part of the pipeline. We supply a simple interface to construct and annotate interactive hierarchical cluster plots, as well as for tree structure visualization using the Interactive Tree of Life (iTOL) web tool ^48^. In particular, to fully leverage the advantage of tree structures for users to identify neighborhoods at any depth of the tree, we implement ways to summarize the tree into clusters (for a more succinct view) that can then be expanded into higher resolution subtrees (**Supp. Fig. 6**). Finally, both HMM and vector based methods for selecting representatives are available to the user.

## Discussion

Protein sequence data has been growing at an unprecedented rate and scalable methods are needed for exploring the sequence space of ever-expanding protein superfamilies. Here, we present three different methods for defining neighborhoods in sequence space, two methods of visualization (including the new hierarchical cluster plot) and two methods for selecting representatives from sequence neighborhoods. We provide an analysis pipeline combining the different methods of neighborhood detection, visualization and representative selection (ProteinClusterTools, https://github.com/johnchen93/ProteinClusterTools). Importantly, we demonstrate the ability for end-users to analyze large sequence spaces on typical end-user desktop hardware, making analyses of large custom datasets more accessible.

Hierarchical visualization methods provide a way to view sequence space in an analogous manner to the organization of sequences (*e.g.*, ranging from distant to close homologs). It also makes it easy to identify subsections of sequence space that can then be focused on for higher resolution separation (*e.g.*, identifying isofunctional families, which can emerge at different cut-offs from each other). The hierarchical cluster plot is the most succinct way to visualize protein sequence space, and is generally applicable with any set of neighborhood definitions; for example, one could base clusterings on pairwise structural similarity or use a different clustering algorithm from those presented. In contrast, the tree based view provides the most detailed resolution of sequence relationships and allows for the most flexible choice of cluster separation, but is limited to hierarchical clustering, which is the least scalable.

We implement three different methods for neighborhood detection, each with its own strengths and weaknesses. The homology based approach uses a well established similarity metric (bitscore, BLOSUM62 matrix) that is interpretable, while the vector representations are not human-interpretable but allow for the application of a broader toolset for dimensionality reduction and clustering. In terms of scalability, the computation time and data size of the homology method increases linearly with the number of sequences, as most homology search tools (BLAST, MMSeqs) will limit the number of hits by the significance of the homology (E-value). There is no such statistical filter as for vector representations at the moment, and any pairwise analyses (*e.g.*, hierarchical clustering) must explore all possible pairwise combinations, making the data expand at an exponential rate. However, methods such as K-means clustering, which do not require all pairwise comparisons to be stored at once, can bypass such a limit. The three neighborhood detection methods are generally in consensus on neighborhood memberships, likely because the similarity metrics of homology and vector distances both correlate with sequence identity. Of course, different substitution matrices for homology or different pLMs for generating vector representations from those applied here could change the outcomes.

The use of the entire available sequence space to generate phylogenetic trees shows the utility of hierarchical clustering for the identification of representative sequences. In the example phylogeny provided here (**Fig. 5**), adding more diverse sequences greatly increased sequence space coverage and at the same time improved tree topology and replicability. While a simpler BLAST based approach will most likely still provide information on separation of major clades, it will miss other sequences that could be important for inferring evolutionary relationships. In particular, the discovery of extra clades when sampling the sequence space more evenly may aid in the formation of evolutionary hypotheses, as well as providing more information for studying evolutionary histories using ancestral sequence reconstruction (ASR)^49–51^. A diverse but even sampling of a neighborhood with known function could also aid in protein engineering, either by providing alternate starting points for engineering function^52^, or by incorporating a broader set of related sequence data to draw from when making designs through coevolution^53^, phylogenies (*e.g.*, ASR)^49,54^ or machine learning models ^55^.

Moving forward, sequence exploration methods are limited by the annotations that the user can analyze, as most sequences are unexplored. The unsupervised clustering methods in this study can provide some structure and topology to the protein families, but the function of unknown sequence space still needs to be illuminated. A limiting factor in sequence space annotation is the lack of experimental characterization, leaving the vast majority of sequence space unexplored and unannotated. This is both a limitation and an exciting opportunity, as unexplored sequence neighborhoods offer the potential for exploring new proteins and enzymes, which can deepen our understanding of biology and provide new proteins for use in biotechnology. In terms of further leveraging existing data, inference by homology or by training of protein language models can be used to try and predict sequence properties to fill the gap in information, but ultimately the accuracy of these predictions will need to be tested. A joint effort to explore sequence space with a global view can help uncover understudied regions that can be further characterized. Focusing on regions of unknown function could enhance the diversity of known functions, and provide better sequence-function data that could be used for protein engineering or function prediction.

## Methods

### Software availability

The hierarchical cluster plot and tree structures were generated and visualized with the pipeline (ProteinClusterTools) described in this paper. ProteinClusterTools can be installed as a python package from the Python Package Index (https://pypi.org/project/ProteinClusterTools/). The source code for the pipeline, including guides on installation and usage, can be found on GitHub (https://github.com/johnchen93/ProteinClusterTools).

### Datasets

The FMN-binding split barrel superfamily (also known as the flavin/deazaflavin oxidoreductases, FDORs) was identified on InterPro using sequence homology searches for representatives in each previously classified FDOR families (A, AA and B families)^16^. We found all representatives fall under FMN-binding split barrel, homologous superfamily (IPR012349) which has ∼323k sequences. We also include the Haem oxygenase HugZ-like superfamily (IPR037119), which has ∼13k sequences (∼6k overlapping with IPR012349). Sequences for both superfamilies were collected from UniProt. For neighbourhood detection, we reduce the redundancy of all collected sequences to just the 100% ID representatives using MMSeqs easy-cluster^56^, resulting in 299,624 sequences.

### Functional Annotations

We retrieved all metadata of the FMN-binding split barrel superfamily from UniProt, collecting all reviewed entries and those with associated structures in the PDB. The reviewed UniProt entries were filtered to exclude annotations based on similarity or UniRule unless they had an associated publication. All remaining entries were manually checked to confirm the functional and co-factor annotations. The list of all PDB IDs in the superfamily were retrieved jointly using those associated with UniProt entries, or from searching the PDB directly using the superfamily codes (IPR012349 and IPR037119), which resulted in a list of nearly fully overlapping structures. Structures from each unique sequence were checked for co-factor and function from the structure as well as any associate publication. Co-factor information from structural alone was tracked, but not used in the final annotations. The final set of annotations combined the UniProt and PDB based information, which was then used to annotate the SSN (**Supplementary Data 1**).

### Homology based neighbourhood detection

For homology based neighbourhood detection, we first construct a conventional SSN by performing all-by-all pairwise alignments using MMSeqs easy-search on default settings (using BLOSUM62 matrix, hits require E-value < 1e-3) but with increased maximum hits (raised from 300 to 1000). The SSN was generated from all 100% ID representatives of the FMN-binding split binding superfamily, resulting in ∼278M pairwise bitscores (edges) without any cut-offs. To decide on the cut-offs, up to 1M pairwise bitscores were sampled to construct a distribution. The cut-offs were then selected based on the 10,25,50,75,90 percentile values of the bitscore distribution. In addition, we conduct a sweep of cut-offs from 100 to 250 in 10 bitscore intervals. We use bitscore as the metric for homology instead of the E-value (which is also common), as the E-value is affected by the total dataset size and lacks resolution at high bitscores (minimum E-value is 1e-180) while bitscores can continue to scale with homolgy.

Neighbourhoods are defined as the isolated clusters at each cut-off (any sequences connected with each other, but not to any other clusters), also known as the “connected components” of the graph. While this could be performed with existing algorithms (such as using NetworkX), it becomes increasingly difficult with larger datasets as the entire network must be loaded at once. Instead, we implement an alternate algorithm that streams the network file in chunks, and builds cluster definitions for each chunk. This allows the whole file to be processed in a manageable size, and allows parallel processing of the entire file, while keeping total RAM usage stable. At the end, clusters from all chunks are merged together. We note that this method of clustering is sensitive to a small number of linkages between clusters (clusters are harder to separate), but is nevertheless the simplest and most scalable that still eventually allows the user to identify isofunctional families. We also note that it is possible to perform hierarchical clustering using the pairwise distance measures such as bitscores, with the key limit for scalability being the ability to ignore data points not in the distance matrix (no significant homology) so as to not construct a full distance matrix (which is exponentially scaling). However, the existing proposed solutions do not have publicly available implementations ^57–59^, and common, easy to use solutions like SciPy require the full distance matrix (which is too large to fit into RAM for our datasets).

### Vector based neighbourhood detection

We convert the amino acid sequences into vector representations by using the ‘extract.py’ script provided with the pLM Evolutionary Scale Modeling (ESM)^17^. Specifically, we use the ESM1b model with 650M parameters (esm1b_t33_650M_UR50S) to encode our sequences. For each embedded sequence the model produces a final hidden state matrix of N x 1280, where N is the protein length; *i.e.*, every amino acid position is converted into a vector of length 1280. We take the mean of all vectors along the sequence dimension (mean-pooling) so that any given sequence has the same final vector length of 1280. For downstream clustering, all sequence vectors are further reduced to 30 dimensions using PCA.

To perform K-means clustering, we use the bisecting K-means implementation from scikit-learn. However, we found that above ∼30k sequences, the clustering at different cut-offs (number of desired clusters, K) no longer formed consistent parent-child relationships across cut-offs. Hence we apply adjustments to the method. Bisecting K-means implemented by scikit-learn produces a “bisecting tree”, that is used to make decisions on which clusters to divide in the next step. This creates a full tree structure, but the final sequence labels in the tree are normally erased to save memory, and so we make a single edit to the code to retain the tree’s information. We start the process for creating a hierarchically consistent set of K-means clusterings, by running this slightly adjusted bisecting K-means (using Euclidean distance) at the maximum desired number of clusters K. Then, we extract the tree structure, where each branch length between a parent cluster and the resulting split children in the tree is the difference of the spread (sum of squared errors) within the parent and child cluster. The tree structure is then converted into a linkage matrix using single linkage method, and flattened into a smaller number of clusters K using the ‘fcluster’ function of SciPy, ensuring the hierarchy is consistent across cut-offs. We scan a range of cut-offs from 3 to 10,000 to determine appropriate cut-offs for visualization.

To perform hierarchical clustering, we use the hierarchy module of the SciPy package. However, the full dataset of ∼300k sequences is too large for hierarchical clustering due to the hierarchical clustering using vectors always constructs the full pairwise distance matrix (N choose 2 combinations, ∼332Gb RAM required for 300k sequences), as there is no significance measure like the E-value that can be used to exclude non-homologs; although most hierarchical clustering implementations will require a full distance matrix and still be limited in scalability regardless. We downsample the data by redundancy reduction and find that the representatives at 70% ID (∼100k sequences) can fit within 64Gb of RAM. To conduct the clustering itself, the linkage matrix is constructed from the vector representations of the 70% ID representatives using the SciPy linkage function (with method=weighted, and metric=cosine). The linkage matrix is ‘flattened’ into clusters using the ‘fcluster’ function by specifying the number of desired clusters (criterion=maxclust). Using the flattened clusters at different cut-offs, we can create a hierarchical clusters plot. Alternatively, we can process the full tree structure generated by hierarchical clustering (converted to Newick format) at a given number of clusters, by simplifying the tree such that each leaf is a cluster, and the child tree of the cluster is extracted for separate analysis. Trees can be annotated using Protein Cluster Tools as a convenient way to apply annotations compatible for interactive visualization with Interactive Tree of Life (iTOL). We scan a range of cut-offs from 3 to 10,000 to determine appropriate cut-offs for visualization.

### Comparison of cluster agreement

We use the implementations for the adjusted Rand index (ARI) and adjusted mutual information (AMI) from scikit-learn. Both metrics are adjusted for random chance so that increasing the number of clusters does lead to higher values unless the agreement also improves. We calculate the agreement between each pair of clustering methods (between homology, K-means and hierarchical clustering) at all pairs of cut-offs at regular intervals. For hierarchical clustering, since the dataset was downsampled to the 70% representatives, we apply the labels of the representatives back to their group members at 100% ID to match the number of data points in the homology and K-means methods.

### Hierarchical cluster plots

Hierarchical cluster plots are constructed with the following approach. At each cut-off, all clusters are filtered to be above a certain size (number of sequences) or to contain a certain sequence. Clusters that pass the filter are grouped by the parent they belong to at a lower cut-off (or all clusters at the lowest level), and packed together using physics simulation using PyBox2D. Clusters are represented as circles, and scaled so the area matches the cluster size. After packing, the child clusters are placed within the parent cluster, based on the center of all clusters. The resulting structure is stored as a layout with size and coordinates for each cluster, which can be visualized with any custom plotting software. The implementation in Protein Cluster Tools uses Bokeh for the interactive visualization, and matplotlib for the figure export.

### Representative selection from neighbourhoods

To select a cluster representative using the HMM approach, all sequences in the cluster are aligned using mafft^60^ with default settings to generate a MSA. The MSA is used to build an HMM using HMMER3 (hmmbuild command), then all sequences in the cluster are searched using the HMM (hmmsearch command). The sequence with the highest score to the HMM is selected as the representative. To select a cluster representative using the vector approach, all vector representations of sequences in the cluster are averaged to generate a centroid vector. The sequence whose vector representation is closest to the centroid (Euclidean distance) is selected as the representative.

### Phylogenetic tree construction

To replicate the previous tree of the FMN-binding split barrel superfamily, we followed the previously published method^45^. We used the 18 structural representatives (PDB IDs: 1USC, 2ECU, 1RZ1, 2IG6, 3R5R, 3H96, 4Y9I, 4YBN, 2HQ7, 2I02, 3DB0, 5BNC, 3TGV, 1VL7, 1RFE, 4ZKY, 4QVB, 3F7E) in a homology search using BLASTP on the current non-redundant protein sequence database with taxa limited to Bacteria or Archaea for the top 500 hits. We excluded search results of 2FQH from the previous study as it was a different fold. All hits had E-values < 1e-9 as previously specified. This resulted in ∼6,000 sequences, which were reduced to 306 representatives (324 including the 18 structural representatives) at 65% sequence identity using MMSeqs easy-cluster with --cov-mode 0 and alignment coverage of 80% (-c 0.8). To determine coverage of BLAST selected representatives in the SSN, MMSeqs easy-search was used to identify the SSN sequence with the highest homology to each representative. A multiple sequence alignment was performed using MAFFT, with the --localpair setting. We removed three sequences from the alignment that had misaligned, with gaps in most of the conserved columns, resulting in a final set of 321 sequences.

To evenly select sequences from the FMN-binding split barrel superfamily, we used cluster tools to first isolate the parent neighbourhoods of the BLAST representatives at the bitscore cut-off of 102, thereby excluding other outlier neighbourhoods. Then, starting at a bitscore cut-off of 150, we sample one representative from each neighbourhood with between 100-1000 sequences through the HMM based representative selection method. Neighborhoods greater than 1000 sequences were further separated at higher bitscore cut-offs in 10 bitscore steps and sampled again, up to a cut-off of 400 bitscore. To exclude potential sequence outliers, we only include representatives of clusters with lengths between 100-300 amino acids, resulting in 346 sequences (364 including the 18 structural representatives). Sequences were aligned using MAFFT with the -- localpair setting.

To construct trees, we curate the MSAs programmatically using CIAlign ^61^ to ensure both datasets are treated with a minimum of human bias. CIAlign was used with the --remove_insertions and --crop_ends option on default settings. The resulting MSAs still had gap heavy regions at the termini, and so each MSA was trimmed to the first conserved block at each termini. Maximum likelihood trees were constructed using the IQ-Tree webserver ^62^, using default model finder and ultrafast bootstrap settings. Each MSA was used to construct 5 tree replicates. All BLAST based replicates had LG+G4 as the best model, while all cluster tools based replicates had LG+F+G4 as the best model. The approximately unbiased (AU) test was performed on each set of replicates, with 10,000 test replicates, resulting in no rejection of topologies in any replicate. We select the tree with the greatest log likelihood as the representative tree for visualization, branch lengths and bootstrap values. Trees were visualized and annotated using Interactive Tree of Life (iTOL) ^48^. Tree topology differences between replicates was calculated as the normalized cluster information distance using the R package TreeDist ^46^.

## Supplementary Info

**Supplementary Figure 1.**
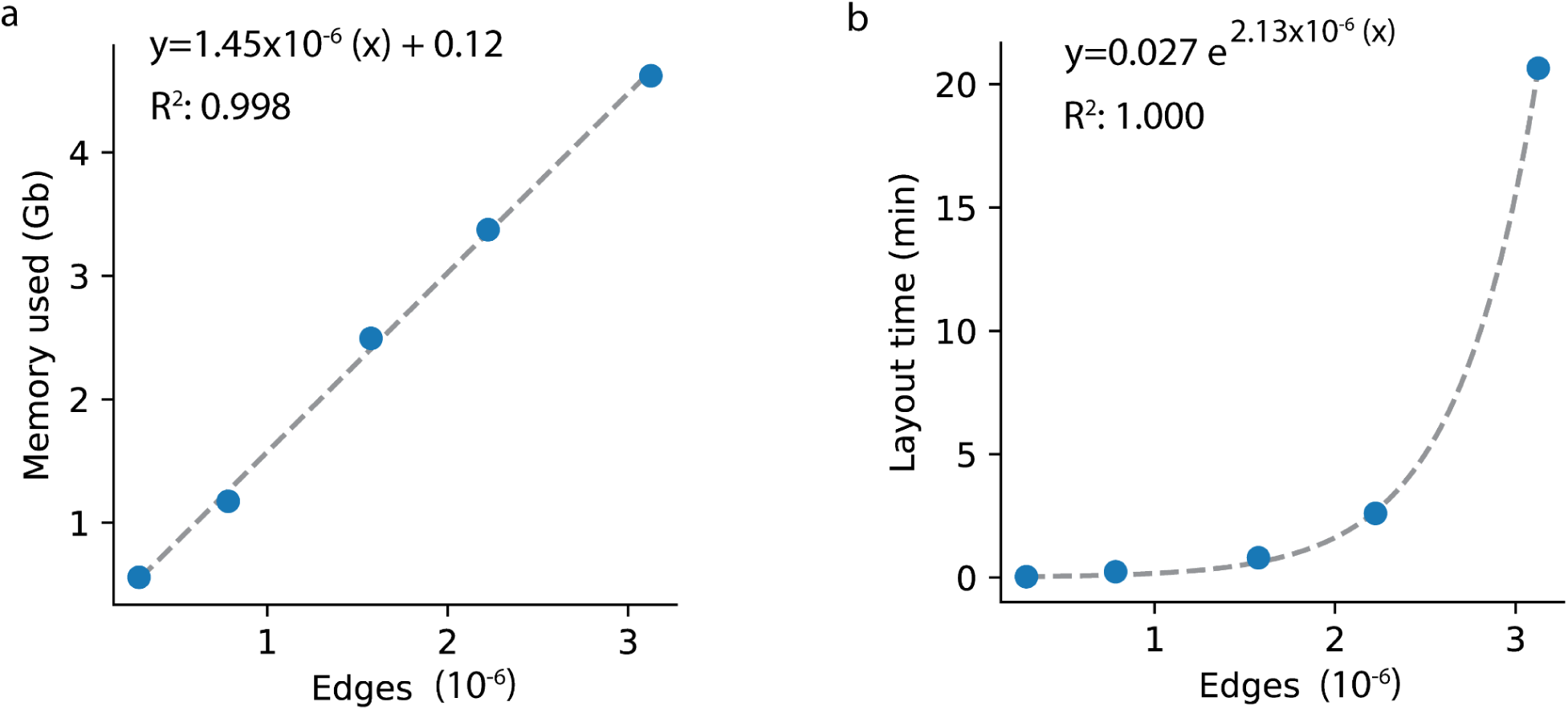
Resource use in Cytoscape as a function of the number of edges in the network. The FMN-binding split barrel superfamily was divided into single networks at bitscore cut-offs of 500-900 in 100 bitscore steps, and time and memory used for visualization in Cytoscape was measured and used to calculate resource use with edge count. (**a**) Gb of memory used for networks of given edge count, fitted to a linear regression. (**b**) Time to generate default layout using Perfuse force directed algorithm for networks of given edge count, fitted to an exponential function. Lines of best fit, equations of the fit and R^2^ are shown in the plot.

**Supplementary Figure 2.**
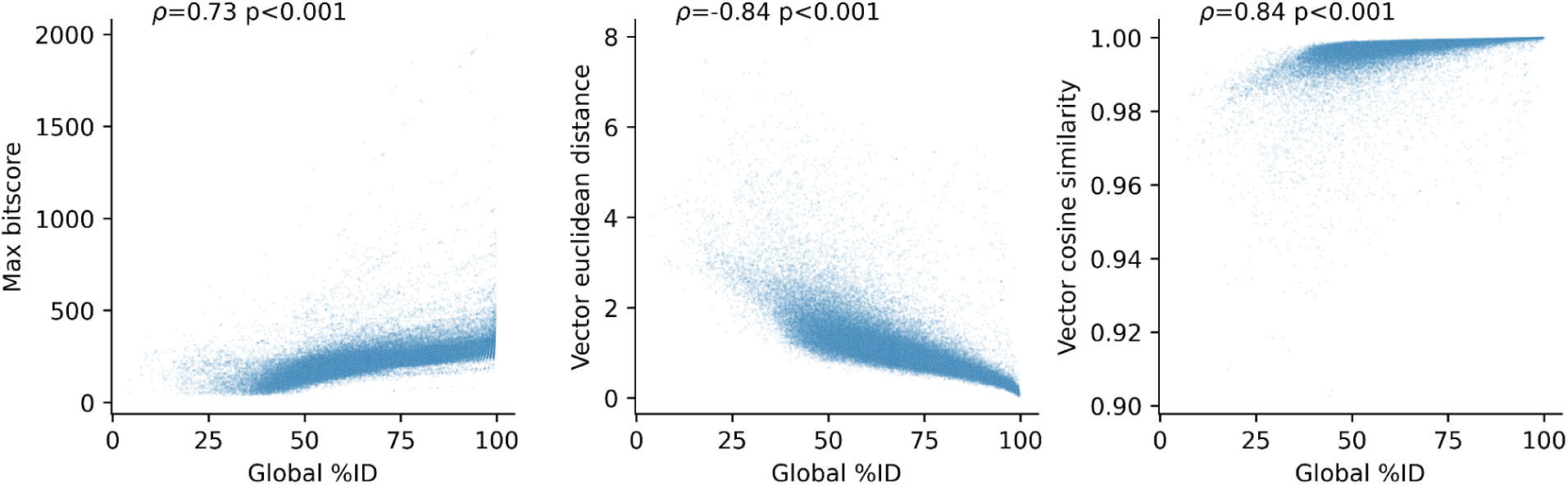
Correlation of sequence similarity metric to global sequence identity. The data was a sample of 71,006 pairs of sequences (excluding self pairing). Max bitscore is calculated from pairwise BLAST between sequences, taking the max value in the case of reciprocal pairs. Vector representations are compared by either their euclidean distance or cosine similarity. The spearman correlation coefficient and p-value are displayed.

**Supplementary Figure 3.**
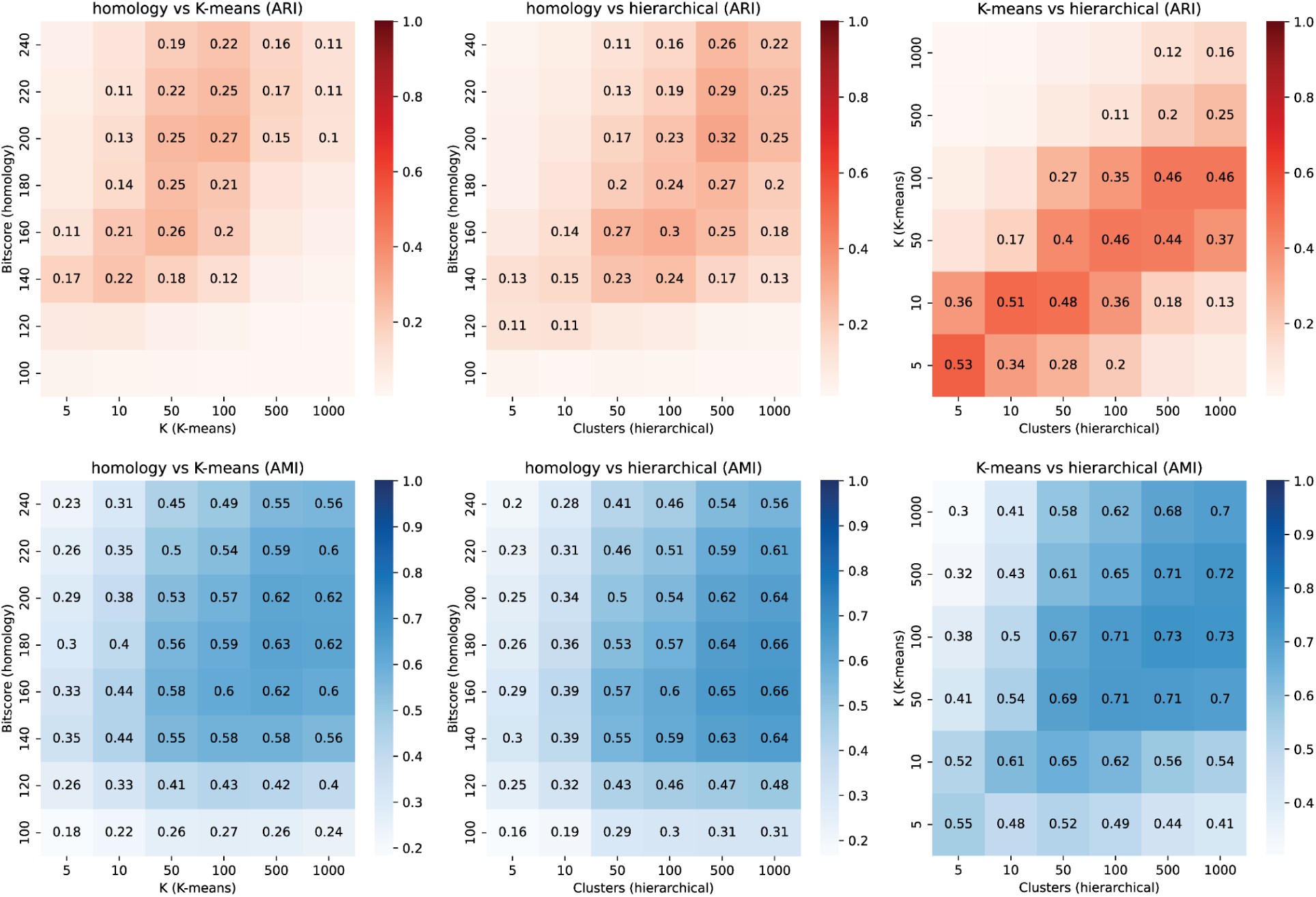
Measure of agreement between cluster assignments between each pair of clustering methods. Heatmaps show the clustering agreement, as measured by the adjusted Rand index (ARI) or the adjusted mutual information (AMI) between each pair of cluster definitions. The clusterings are arranged by the bitscore cut-off for homology, while K-means and hierarchical clustering are arranged by the number of target clusters.

**Supplementary Figure 4.**
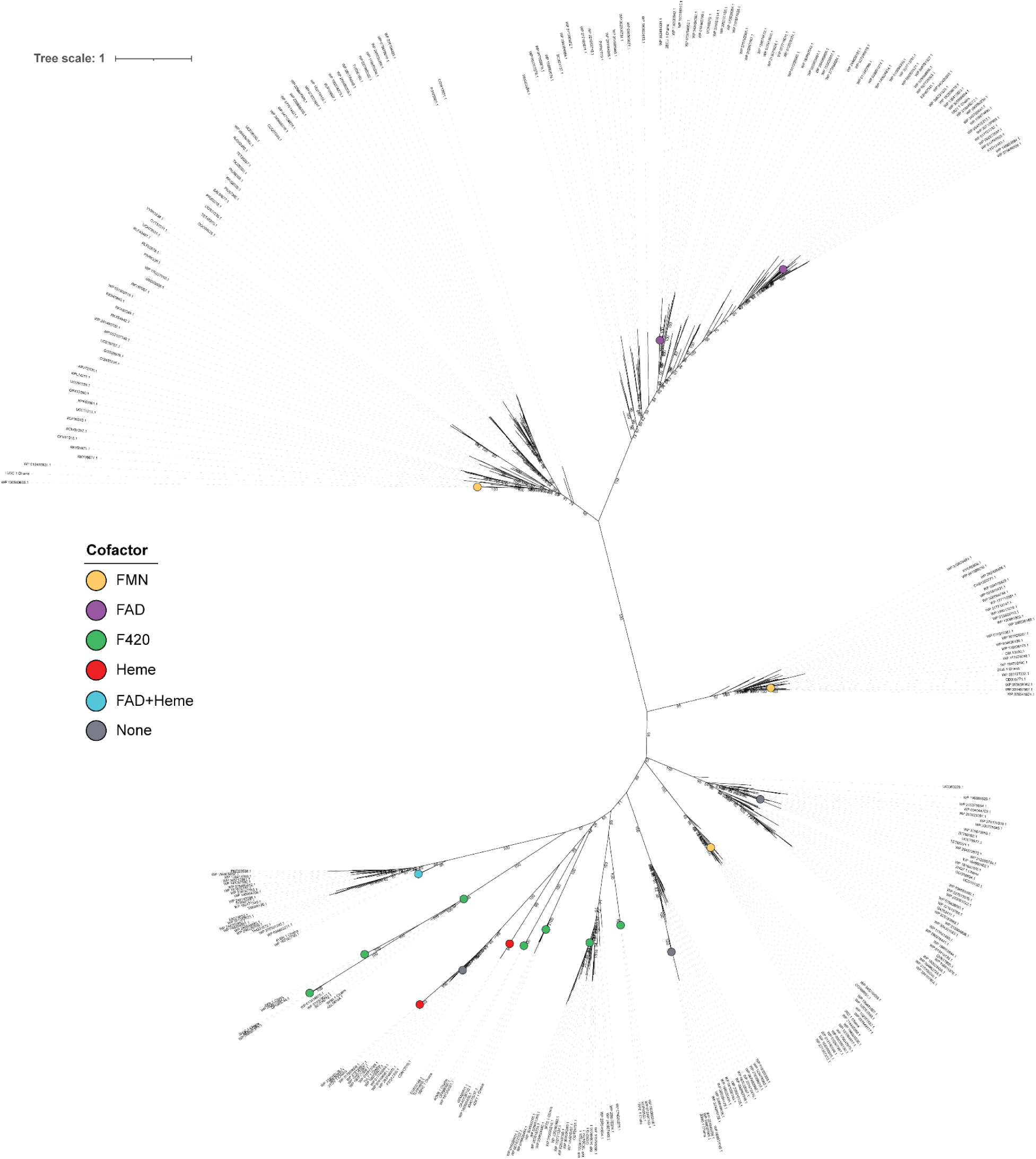
BLAST sampling based maximum likelihood tree of FMN-binding split barrel superfamily. The tree contains 321 leaves. The 18 structural representatives and their co-factors are highlighted. Bootstrap values are displayed as numbers.

**Supplementary Figure 5.**
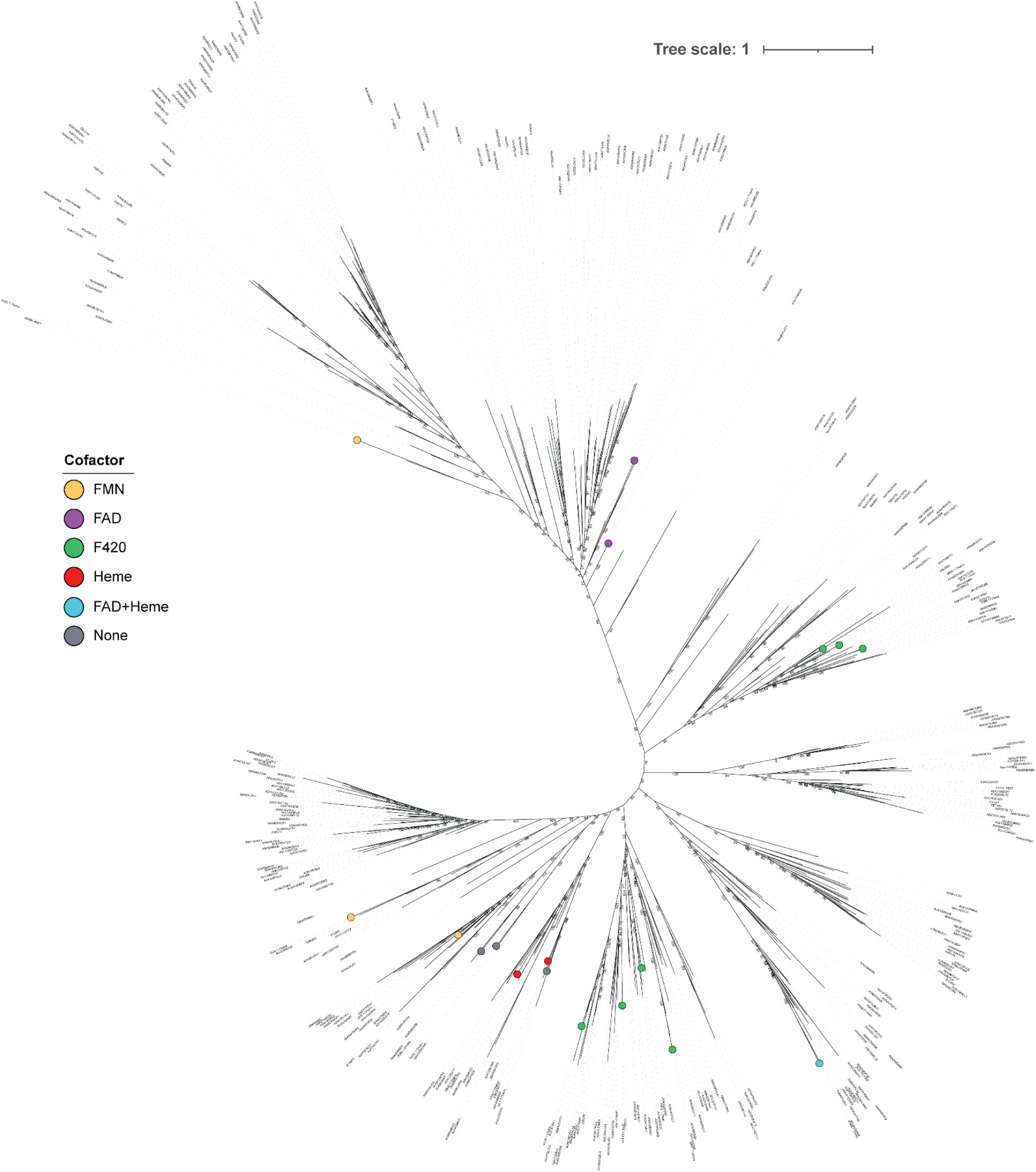
Cluster tools sampling based maximum likelihood tree of FMN-binding split barrel superfamily. The tree contains 364 leaves. The 18 structural representatives and their co-factors are highlighted. Bootstrap values are displayed as numbers.

**Supplementary figure 6.**
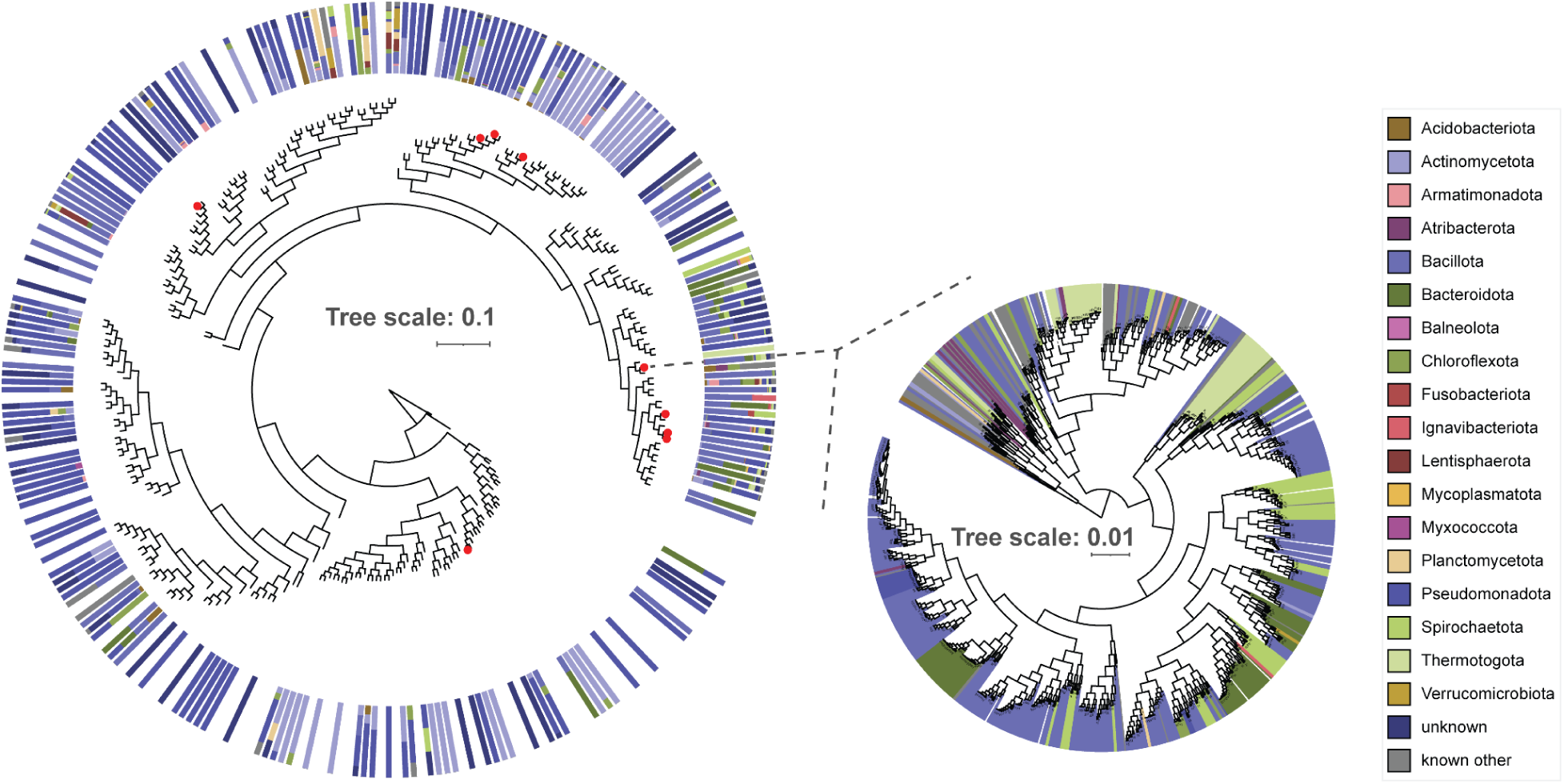
Tree structure visualization of hierarchical clustering results. The tree structure of the Periplasmic binding protein/LacI sugar binding domain (IPR001761). The full tree structure of ∼40k sequences was constructed using the hierarchical clustering method on the sequence vector representations. The full tree was condensed to 300 clusters (left) and annotated according to taxonomy, leaves highlighted in red have SwissProt annotations. The smaller tree on the right illustrates the annotated subtree from one of the clusters.

**Supplementary Table 1.** Observed resource usage for network visualization in Cytoscape.

| **Bitscore** | **Edges (10<sup>6</sup>)** | **Memory (Gb) <sup>a</sup>** | **Layout Time (min) <sup>b</sup>** | **Connected sequences <sup>c</sup>** | **Percent connected <sup>d</sup>** |
| --- | --- | --- | --- | --- | --- |
| 900 | 0.3 | 0.6 | 0.03 | 6,001 | 2% |
| 800 | 0.8 | 1.2 | 0.2 | 7,209 | 2% |
| 700 | 1.6 | 2.5 | 0.8 | 8,838 | 3% |
| 600 | 2.2 | 3.4 | 2.6 | 13,201 | 4% |
| 500 | 3.1 | 4.6 | 21 | 25,837 | 9% |

- a. Memory usage as reported by Cytoscape after layout is completed and unused memory is manually freed. - b. Time for Cytoscape to generate the default layout using Perfuse force directed algorithm. - c. Number of unique sequences connected to any other non-self sequence. - d. Connected sequences as a percentage of total number of sequences (299,624).

**Supplementary Table 2.**
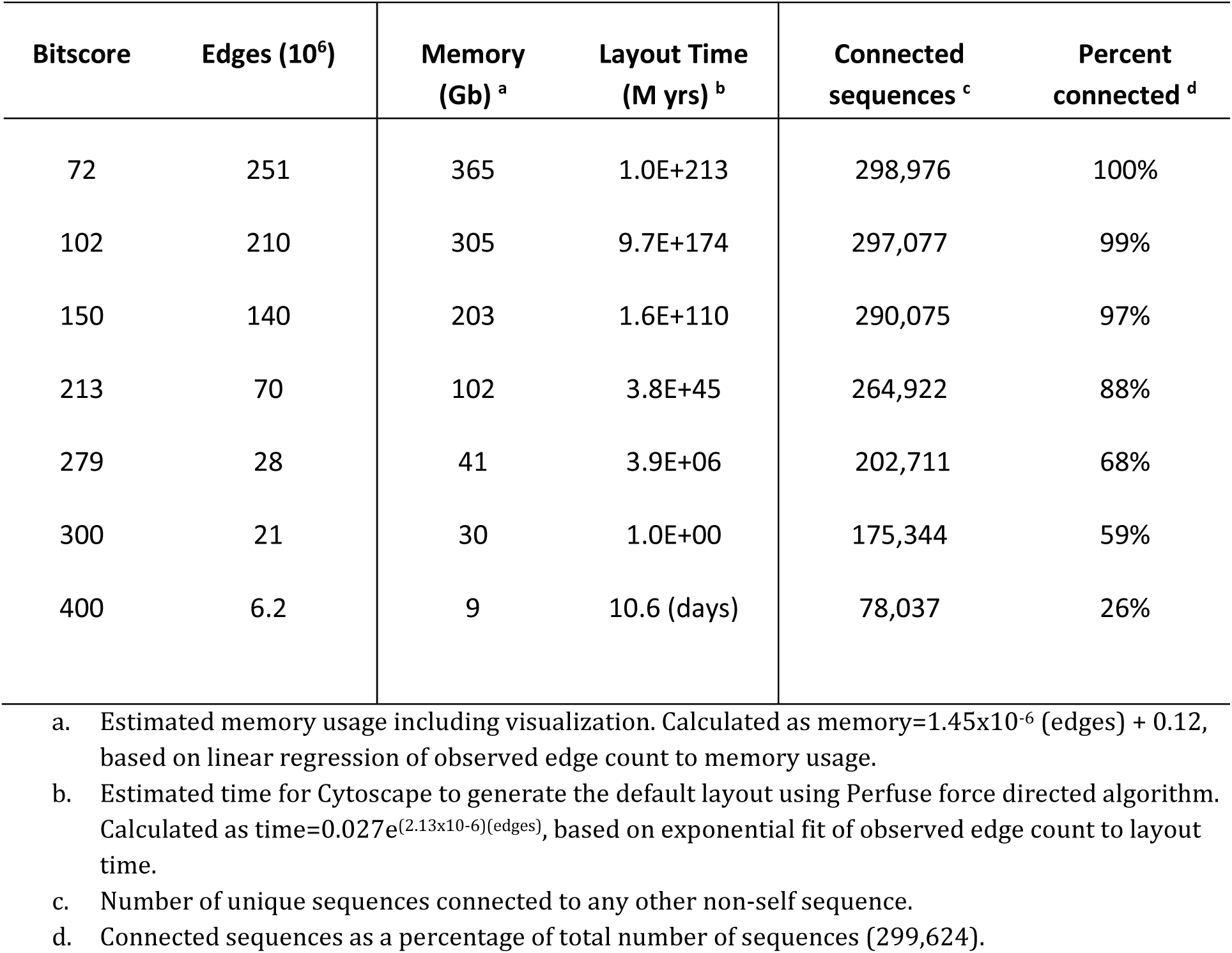
Estimated resource usage in Cytoscape for network of a given size.

